# Extraction and characterization of cellulose from invasive weeds from central Nepal: A potential prospect of environmental management

**DOI:** 10.1101/2022.09.08.507090

**Authors:** Samyog Dhakal, Achyut Tiwari, Archana Adhikari, Shyam Kumar Shrestha, Bikash Adhikari

**Author notes:** Authors with equal contribution (Samyog Dhakal, Achyut Tiwari).

## Abstract

The cellulose is made up of long chains of polysaccharide of glucose molecules. Microfibrils are formed when numerous hydrogen-bonded cellulose chains unite, they are extremely stiff and contribute to physical stability, due to this general ability of forming these microfibrils to form long chains, cellulose is an ideal molecule for the manufacturing of packaging materials and bioplastics. On the other hand, Invasive plant species are one of the major constituents for environmental degradation and its application seems outmost. The main purpose of this study is to extract and identify the composition of cellulose fiber and characterize the fiber of invasive plant species that could be used as a replacement for plastics and textiles in some cases. In this study, Cellulose was isolated from 6 invasive species collected in Nepal’s Ramechhap district using various techniques, the composition of the fiber was identified using AOAC method 973.18, ASTM method D1106-96 and ASTM method E1755-01 and characterized using FTIR spectroscopy with weight analyses. Acid hydrolysis, chlorination, alkaline extraction, and bleaching were among the chemical methods adopted. In all of the samples, there were two primary absorbance peaks. The first occurred at low wavelengths in the 700^−1^,800 cm^−1^ range, while the second occurred at higher wavelengths in the 2,700–3,500 cm^−1^ range.The percentage of lignin within the final sample was determined in the range of 4.4-3.1% and the percentage yield of cellulose was determined within the range of 78-62%.The study shows that the cellulose can be extracted from the taken invasive plant species and can be used for further applications.

## Introduction

Biological invasion by exotic plant species poses a major threat to native plant communities, and alters ecosystem structure and function. Plant invasion is now considered as one of the major causes of biodiversity loss, resulting in significant vegetation change and environmental degradation globally [26, 14, 5].Invasive plant species offer an excitingly cheaper and widely available renewable source of cellulosic fibers [21, 25] and can be used as a tool of controlling highly pervasive use of non-biodegradable plastics which have been creating environmental problems including health hazards.Plastic pollution has been very intense and a global environmental problem because of its wide use in transportation, food, clothing, food industry, product packaging, medicine and agricultural fields has been observed in recent years [4, 26]. Plastics are complex polymers with long repeating chains of molecules and are derived from petrochemicals, they are lightweight, cheaper, highly durable and are of high strength leading to their increasing demand. The emissions of plastic are increasing day by day and will continue to do so despite great efforts of plastic waste reduction, and the global emissions of plastic ranges from 9 to 25 million metric tons per year to the terrestrial and aquatic environment [3, 12].

Invasive alien plant species (IAPS)include alien plant species which become established in natural or semi-natural ecosystems or habitats, and are an agent of change threatening native biological diversity [7]. Rapidly changing climate is expected to increase the frequency and intensity of biological invasions including the invasion of IAPS [21]. It is seen that drought has the ability to enhance the diversity and abundance of invasive plant species in a number of habitats, which is typically mediated by the presence of disturbances [15]. Climate warming has been very much significant in different parts of central Himalaya including the sample collection area of this research (Manthali, Ramechhap: Bagmati Province), known to be one of the drought prone areas in Nepal [24, 17]. There has been higher invasion of IAPS along agricultural land, threatening soil fertility, crop productivity and the existence of native plant species [24].

Both, the plastic pollution (non-biodegradable property) and IAPS are very harmful to our environment which are disrupting natural communities and ecological processes [17] thereby degrading the ecosystem services. Himalayan region is experiencing drought episodes associated with rapid warming of temperatures and increasing uncertainties in rainfall patterns [6]. And it is seen that drought has the ability to enhance the diversity and abundance of IAPS in a number of habitats, which is typically mediated by the presence of disturbance factors [24]. Hence human disturbance, increasing drought and invasion of IAPS can be impactful to the ecosystem, and the management of IAPS has been highly critical for securing agricultural productivity and sustainable management of biological resources.

Researchers have initiated the idea of extracting the cellulose from different plant species checking its quality and quantity, which would play an important role in plastic waste management and also enhancing applicability of invasive plants. Cellulose can be used as the raw material for bioplastics [11]. Cellulose is a linear glucose polymer with alpha-1,4 linkages that is generally organized into microcrystalline forms and is difficult to dissolve or hydrolyze [8] Similar to reinforced concrete, cellulose microfibrils are embedded in an amorphous matrix [13] that provide strength and stiffness to the cell wall by acting as structural elements. Because of its unique qualities, structure, and capabilities, cellulose is one of the most common and widely utilized polymers in the food packaging business[28]. It contributes to the reduction of synthetic packaging and trash. It is low in weight and helps to substantially minimize packing material weight [16]. Cellulose also gives the packing material barrier qualities, decreasing the flow and migration of moisture, lipids, gasses, and solutes [29]. The extraction of cellulose from IAPScould be highly effective to create bioplastic products, and could be one of the most important control solutions for both controlling plant invasion and plastic pollution in the environment.Therefore we sought to extract the cellulose from six different IAPS (*Mentha Canadensis, Chromolaena odorata, Amaranthus spinosus,Eichhornia crassipesXanthium strumarium* and *Biden pilosa*) that are widely distributed in Nepal,in order to explore the possible application of IAPS as the source of cellulose microfibrils. Further, we compared the amounts of extracted cellulose from these plant species.

## Materials and Methods

### Materials

We have collected six IAPS from Manthali, Ramechhap of central Nepal Following IUCN inventory [7] (Figure 1). The species list and related information are given in Table 1. The outer part from the stem was collected and processed for further extraction. Instruments like Crucible, Oven, Desiccators, Autoclave, Soxhlet apparatus, Water Bath, Nylon filter/Sieve (40 mesh), Furnace, FTIR spectrophotometer were used in different time according to the necessity. Similarly, Chemicals like formic acid, sodium hydroxide, Hydrogen peroxide, Ethanol, Sodium Hypochlorite, sodium chlorite, acetic acid, sulfuric acid, Ethylene-Toluene solution were used for laboratory treatment.Acid Detergent Fiber (ADF) according to AOAC method 973.18. Lignin in the fibers was determined as Klason lignin according to ASTM method D1106-96 and ASTM method E1755-01 was used to determine the ash content in the fibers[1].

**Figure 1:**
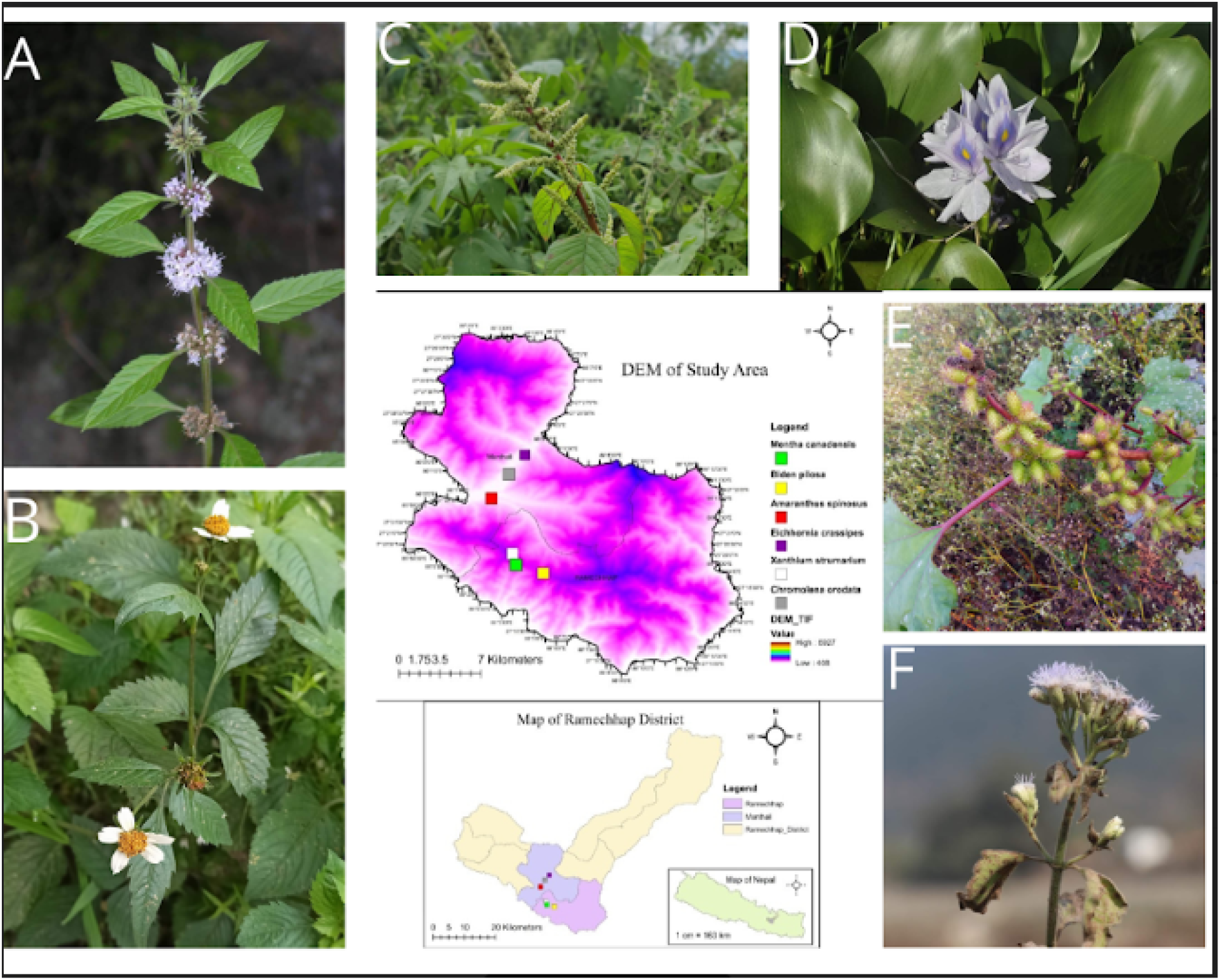
Map of sample collection area in central Nepal, A:*Mentha canadensis* B:*Bidens pilosa* C: *Amaranthus spinusos* D: *Eichornea crassipes* E: *Xanthium strumarium* F: *Chromolena odorata*

**Table 1:** Samples that were collected from the field with the help of inventory [7].

| S.N | Species | Family | Distribution range in Nepal | Native range |
| --- | --- | --- | --- | --- |
| 1 | *Mentha canadensis* L. | Lamiaceae | 200-2500 m | Europe, Asia |
| 2 | *Chromolaena odorata*(Spreng.)<br>King and Robinson | Asteraceae | 850-2200 m | Mexico, C & S America |
| 3 | *Amaranthus spinosus*L. | Amaranthaceae | 150-1200 m | Tropical America |
| 4 | *Eichhornia crassipes* (Mart.) Solms | Pontederiaceae | 200-1500 m | Mexico, C & S America |
| 5 | *Xanthium strumarium*L. | Asteraceae | 100-2500 m | America |
| 6 | *Biden pilosa*L. | Asteraceae | 100-3000 m | Tropical America |

### Fiber extraction

We noticed that the stems of the collected invasive plant species were sensitive to the circumstances of extraction. Strong alkaline environment and/or heating the stalks above 80 °C caused the bark to break down into tiny fibers that were unsuitable for high-value fibrous applications. Based on the yield, length, and strength of the produced fibers, the most ideal conditions for fiber extraction were found after multiple attempts. The peeled fiber was immersed in 0.5 N sodium hydroxide solution with a solution to fiber ratio of 10:1 at room temperature overnight under the ideal circumstances. The fibers were then cleaned multiple times with distilled water before being dried in an oven at 80 Degrees Celsius for 24 hours(Figure 2). They were then sliced to a length of around 5–10 mm. Finally, the wax was removed by boiling for 6 hours in a combination of toluene and ethanol (2:1 volume/volume) in a soxhlet. After that, the fibers were filtered, rinsed in ethanol for 30 minutes, and dried.

**Figure 2:**
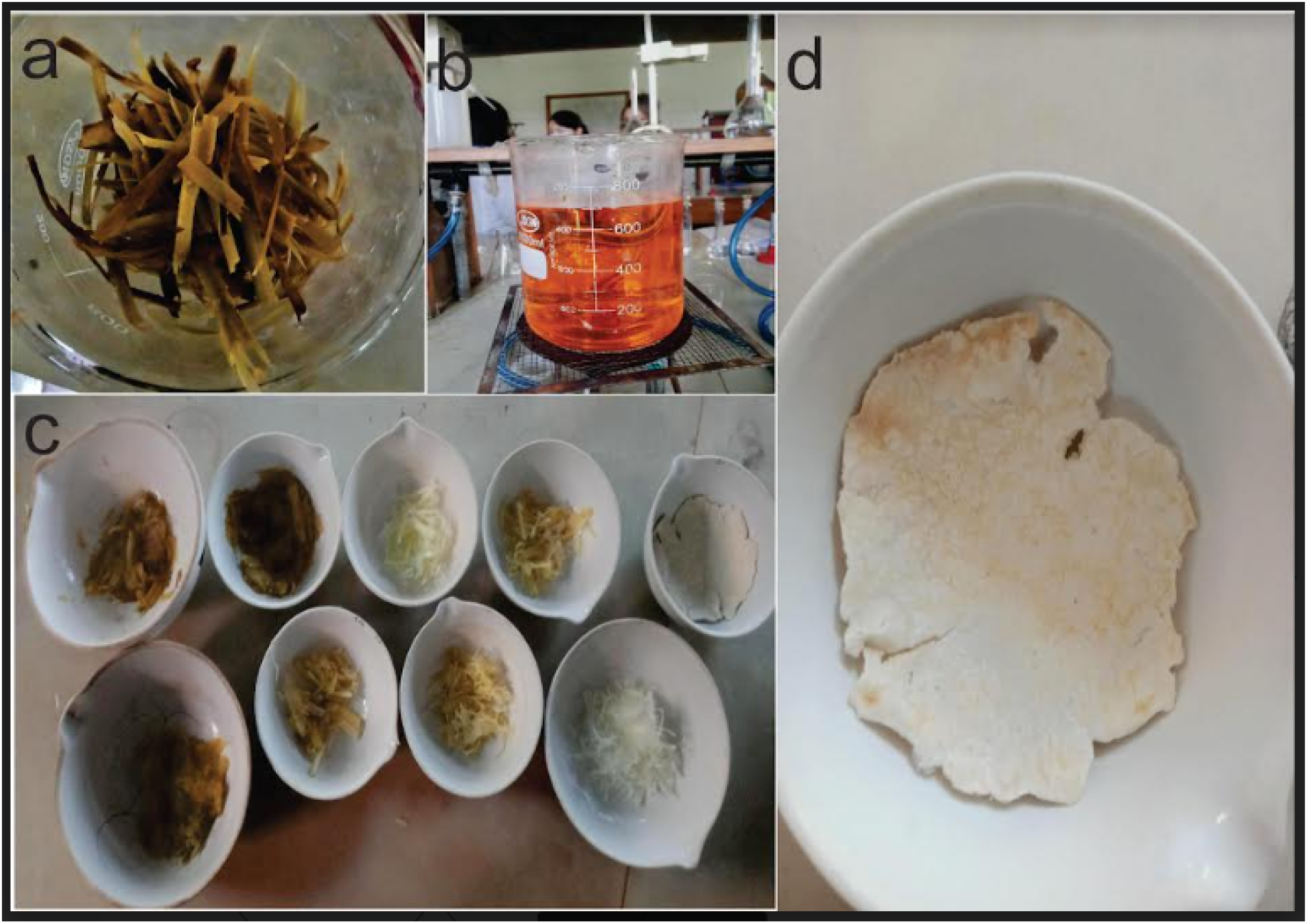
Plant material and extracted cellulose, a: Ground samples before any chemical treatment, b: Treatment with sodium chlorite and boiling after the treatment with hydrogen peroxide, c: Treated ground materials in different phases(White colored samples are in the final phase and ready for FTIR analysis, d: Final Sample ready for FTIR analysis.

**Figure 3:**
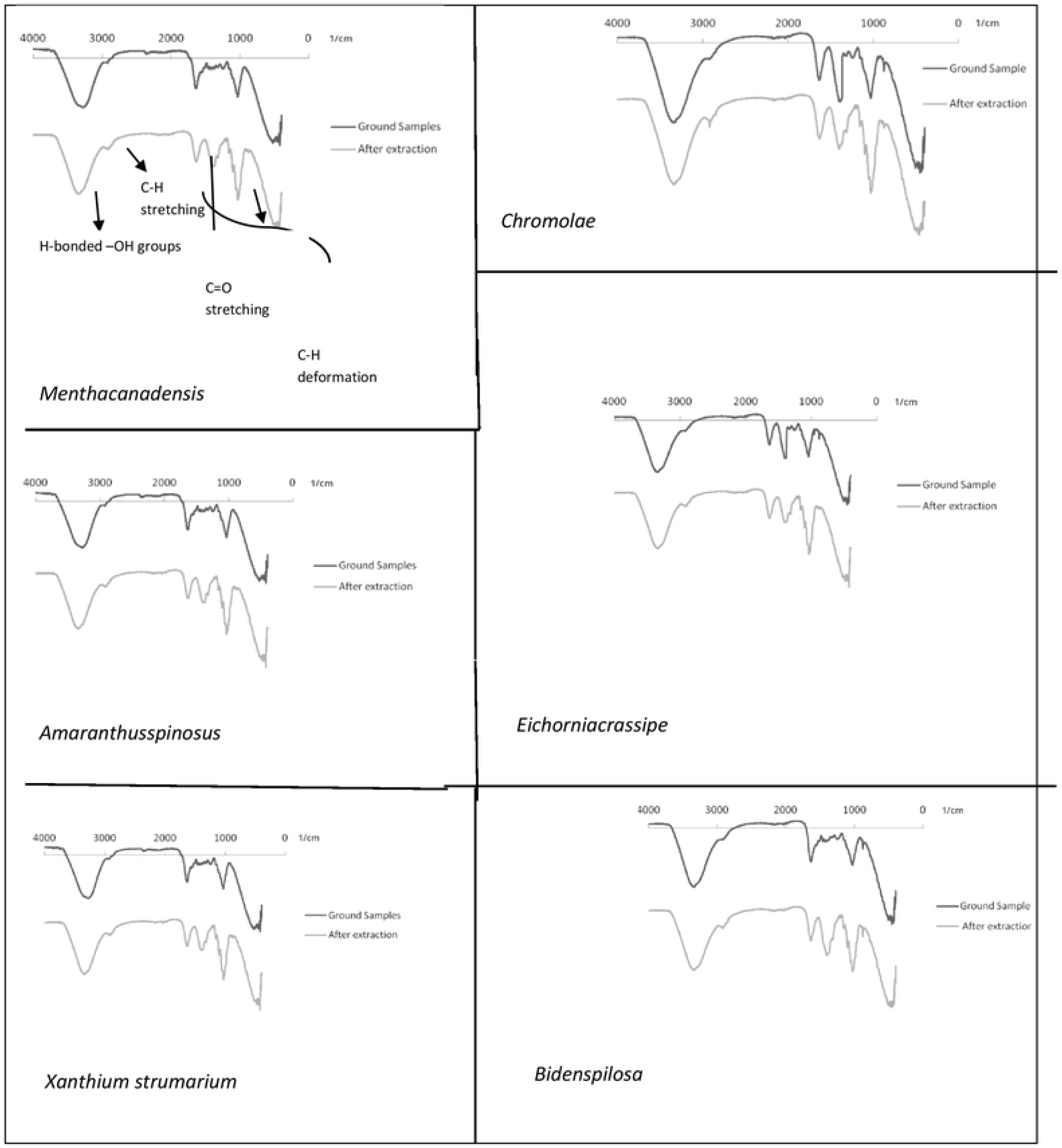
FTIR spectral images of the ground samples and the extracted samples from different invasive plants

### Fiber Composition

Using accepted test procedures, the percentages of cellulose, lignin, and ash in the milkweed stem fibers were identified. AOAC method 973.18 was used to identify the cellulose in the fibers as the Acid Detergent Fiber (ADF) [9]. According to ASTM method D1106-96, the lignin in the fibers was identified as Klason lignin, and ASTM method E1755-01 was utilized to evaluate the fibers’ ash content [1]. Each component was determined using three replications, and the average and the standard deviation are given.

### Lab treatments

i. Pre-treatment with 0.1 M NaOH in 50% ethanol at 45°C for 3 hours with continuous agitation;
ii. Treatment with hydrogen peroxide at pH 11.5 (buffer solution) and 45°C, 3.0 percent H_2_O_2_ for 3 hours each under continuous agitation;
iii. Treatment with 0.7 w/v sodium chlorite NaClO_2_ : holocellulose (a-cellulose + hemicellulose) generation by progressive lignin removal; at pH 4 (buffer solution) boiling for 2 hours with a fiber to liquor ratio of 1:50 and treatment with sodium bisulphate solution 5% w/v;
iv. Holocellulose treatment with 17.5 w/v NaOH solution;
v. Filtering, washing with distilled water and washing with 95 percent ethanol; washing with water and washing with 95 percent ethanol again; drying at 60°C in a vacuum oven until constant weight.
vi. Fiber composition was determined using above mentioned standard procedures.

Ground Sample percentage reduction for cellulose extraction was calculated using the formula:

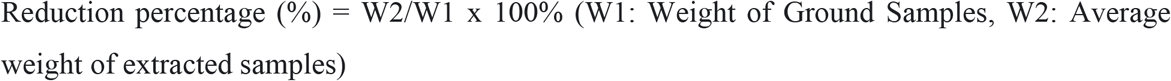

### Characterization

FTIR Spectrometer was used to analyze the Chemical bonding present in newly made cellulose from fiber, raw fiber. The specific chemical groups present in the sample will be determined using spectrum data in automated spectroscopic software based on the infrared absorption frequency range 600–4000 cm^−1^ [22], using IRPrestige-21” present in the laboratory of the Department of Pharmacy, Kathmandu University, Dhulikhel, Nepal. The initial ground samples and final obtained samples were characterized using this method at different time periods.

## Results and Discussions

The FTIR analysis was conducted to identify the presence of different functional groups in the samples. Based on FTIR analysis, there have been two main absorbance peaks in all of the samples. The first at low wavelengths in the 700–1,800 cm^−1^ range, and the second at higher wavelengths in the 2,700–3500 cm^−1^ range, respectively. For each component, however, distinct absorption peaks could be found.

Lignin revealed distinct peaks in the 1,500–1,600 cm^−1^ range, which corresponded to aromatic skeletal vibrations. Furthermore, absorbance in the range of 1,830 cm^−1^ to 1,730 cm^−1^ was detected due to the presence of functional groups such as methoxy –O–CH3, C–O–C, and aromatic C=C. After the same thorough drying process, all of the FTIR spectra were generated. The water adsorbed in the cellulose molecules is exceedingly difficult to remove due to the cellulose-water interaction.

### Composition Analysis

Based on the experimental method, 15 gm of the sample were taken for the experiment and the reduced sample after the final processing of cellulose extraction were obtained as 3.2 - 5.6 gm among the plants (Table 2)

**Table 2:** The reduced sample after the final processing of cellulose extraction.

| S.N | Species | Final weight of the sample (gm) |
| --- | --- | --- |
| 1 | *Mentha canadensis* | 4.2±0.07 |
| 2 | *Chromolaena odorata* | 3.7±0.15 |
| 3 | *Amaranthus spinosus* | 3.2±0.12 |
| 4 | *Eichhornia crassipes* | 5.1±0.17 |
| 5 | *Xanthium strumarium* | 5.6±0.16 |
| 6 | *Biden pilosa* | 4.3±0.22 |

Fiber composition of all the samples were examined, each component was determined using three replications. The highest amount yield of Cellulose is seen in *Chromolaena odorata*(72%) while the lowest is seen in *Xanthium strumarium*(62%). The comparative details on the average with standard deviation are given in Table 3.

**Table 3:** The reduced sample after the final processing of cellulose extraction.

| S.N | Species | Cellulose (%) | Lignin (%) | Ash (%) |
| --- | --- | --- | --- | --- |
| 1 | *Mentha Canadensis* | 72±1.2 | 4.4±0.2 | 1.9±0.02 |
| 2 | *Chromolaena odorata* | 75.3±1.3 | 4.2±0.8 | 2.1±0.03 |
| 3 | *Amaranthus spinosus* | 78.7±1.2 | 3.6±0.7 | 2.4±0.03 |
| 4 | *Eichhornia crassipes* | 66±1.7 | 3.1±0.1 | 1.1±0.02 |
| 5 | *Xanthium strumarium* | 62.7±1.3 | 4.1±0.3 | 1.4±0.05 |
| 6 | *Biden pilosa* | 71.4±1.2 | 3.9±0.9 | 2.1±0.04 |

Our results showed that different invasive weeds’ cellulose contents varied between 72 and 62 percent (Table 3). The yield of cellulose from delignified areca fibers was 65% (w/w) on a dry weight basis in a study conducted by [18]. Similar results were obtained by [23] in their study on *Calotropis procera* fiber (CPF), where the cellulose concentration was determined to be 91.3 wt% upon bleaching.

### Structural Characterization

We generated all the FTIR spectra after the same thorough drying process. The water absorbed in the cellulose molecules is exceedingly difficult to remove due to the cellulose-water interaction [2].The FTIR spectra of the ground sample and extracted cellulose of all the taken invasive plant species are given below. In all of our samples, the Broad bands are seen at around 3400 cm^−1^ and 2890cm^−1^ which relate to stretching of H-bonded −OH groups and C-H stretching absorption of methyl and methylene units. The band at around 3400 cm^−1^ seems to be slightly narrower in the extracted sample, which signifies the higher number of OH-groups. The Similar narrower band was observed in the extracted sample at 3332cm^−1^ of H-bonded OH groups and at 2901 cm^−1^ to the C-H stretching on the work done by[10], in *Retama retama* stems. The cellulose extraction from rice straw waste showed that the broad band was seen between 3600-3100 cm^−1^ and 2912 cm^−1^ which is associated to O-H stretching of hydrogen bond and CH_2_ stretching vibration respectively; these bands show the distinguishing features of cellulose which was seen in this work as well [19]. Similarly of cellulose characterization of *Calotropis procera on* the CPF, peaks at 1163 cm^−1^ and 899 cm^−1^ were related to hemicellulose’s C-O stretching and C-H vibrations, respectively [23]. In our study, we have obtained similar peaks at around 1157 cm^−1^ and 901 cm^−1^ were observed. The presence of such peaks in both works’ samples proved that lignin and hemicellulose were present. Also in the de-lignined CPF, the absorption band at 1732 cm^−1^ and 1502 cm^−1^ disappeared, indicating that the lignin was effectively removed. The band at 899 cm^−1^ vanished from the bleached fiber, confirming that the hemicellulose had been removed. In our research as well, the absorption bands at 1736 cm^−1^ and 1516 cm^−1^ exhibited a decrease or removal in treated samples, indicating that the majority of the hemicelluloses and lignin was removed. The aromatic C=C ring stretching has been correlated to the absorption bands at 1604 cm^−1^ and 1516 cm^−1^. While the –CH_2_ scissoring, C-H asymmetric, –OH bending, C-O antisymmetric, C-O-C pyranose ring skeleton are represented by reaming bands at 1419 cm^−1^, 1372 cm^−1^, 1312 cm^−1^, 1157 cm^−1^ and 1026 cm^−1^, correspondingly. The adsorbed water is responsible for the absorption band at 1628 cm^−1^ seen in the spectra.

## Conclusion

This study confirms the removal of hemicelluloses and lignin validated by FTIR spectra, which were based on the typical peaks of lignocellulosic components and also signifies the processed sample as the cellulose.The weight analysis of different species shows the reduction of weight of the ground sample, the reduced weight was of lignin and hemicelluloses which was confirmed after FTIR analysis. The percentage of lignin within the final sample was determined within the range of 4.4-3.1% and the percentage yield of cellulose was determined within the range of 78-62%. The weight of the final sample was determined within the range of 5.6 to 3.2 gm. The study shows that the cellulose can be extracted from the taken invasive plant species and can be used for further applications. The study provides space for further study on cellulose of invasive plant species using XRD, SEM and other analysis methods that will increase the validity of the study. Further studies could be conducted on different invasive plant species as the amount of extract may differ in plants as per their development stage. Even after drying, the absorbed water doesn’t seem to be reduced significantly, the study reflects the necessity of better chemical processes in upcoming studies that can reduce the amount of water absorbed in the sample. The result from the study can be a source to the policy maker that would help them to make policy regarding the alternative of plastics and management of invasive plant species.

## Reference

1. American Society for Testing and Materials. (1970). Annual book of ASTM standards. Philadelphia, Pa: ASTM.

2. Baird, S. M., John D., Hamlin, O’Sullivan.A., Whiting.A., (2008), An insight into the mechanism of the cellulose dyeing process: Molecular modeling and simulations of cellulose and its interactions with water, urea, aromatic azo-dyes and aryl ammonium compounds, Dyes and Pigments, Volume 76, Issue 2, Pages 406–416, ISSN 0143-7208, https://doi.org/10.1016/j.dyepig.2006.09.01.

3. Borrelle, S. B. et al. (2020), Predicted growth in plastic waste exceeds efforts to mitigate plastic pollution. Science 369, 1515–1518 (2020).

4. Carpenter, E. J., Anderson, S. J., Harvey, G. R., Miklas, H. P., Peck, B. B. (1972), Polystyrene spherules in coastal waters. Science 178, 749–750 (1972).

5. Charles H, Dukes JS, 2007. Impacts of invasive species on ecosystem services. In: Nentwig W (Ed.), Biological Invasions. Springer, Heidelberg, p.217–237. https://doi.org/10.1007/978-3-540-36920-2_13

6. IPCC, 2014. Climate Change 2014: Impacts, Adaptation, and Vulnerability. Part A: Global and Sectoral Aspects. Contribution of the working group II to the fifth assessment report of the Intergovernmental Panel on Climate Change. Cambridge University Press, Cambridge.

7. IUCN(2005). An inventory and assessment of invasive alien plant species of Nepal. IUCN.

8. Jeffries, T. W. 1990. Biodegradation of lignin-carbohydrate complexes. Biodegradation 1, 163–176, (1990).

9. K. Helrich, Official Methods of Analysis, Association of Official Analytical Chemists, Virginia, 170 (1990).

10. Khenblouche, Abdelkader & Bechki, Djamel & 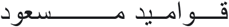, Gouamid Messaoud & Charradi, Khaled & Segni, Ladjel & Hadjadj, Mohamed & Boughali, Slimane. (2019). Extraction and characterization of cellulose microfibers from Retama raetam stems. Polímeros. 29. 10.1590/0104-1428.05218.

11. Isroi, A Cifriadi, T Panji, Nendyo A Wibowo, K Syamsu (2017).Bioplastic production from cellulose of oil palm empty fruit bunch, IOP Conference Series: Earth and Environmental Science, (65), 012011.

12. LauW. W. Y. et al. (2020),Evaluating scenarios toward zero plastic pollution. Science 369, 1455–1461 (2020).

13. Leonowicz, A., Cho, N. S., Wasilewska, W., Rogalski, M., & Luterek, J. (1997). Enzymes of white rot fungi cooperate in biodeterioration of lignin barrier. Journal of The Korean Wood Science and Technology. 25, 1–20.

14. Mack, R. N., Simberloff, D., Mark Lonsdale, W., Evans, H., Clout, M., & Bazzaz, F. A. (2000). Biotic invasions: causes, epidemiology, global consequences, and control. Ecological applications, 10(3), 689–710.

15. Mojzes, A., Ónodi, G., Lhotsky, B. et al. Experimental drought indirectly enhances the individual performance and the abundance of an invasive annual weed. Oecologia 193, 571–581 (2020).

16. Pooresmaeil, M., Nia, S. B., & Namazi, H. J. (2019). Green encapsulation of LDH (Zn/Al)-5-Fu with carboxymethyl cellulose biopolymer; new nanovehicle for oral colorectal cancer treatment. International journal of biological macromolecules, 139, 994–1001 (2019).

17. Rai, K.P., Singh, J.S. (2020) Invasive alien plant species: Their impact on environment, ecosystem services and human health, Ecological Indicators, 111, 106020 (2020).

18. Ranganagowda R. P. G, Kamath S. S, Bennehalli B. Extraction and characterization of cellulose from natural areca fiber. Mat. Sci. Res. India; 16 (1).

19. Razali, N.A.M.; Mohd Sohaimi, R.; Othman, R.N.I.R.; Abdullah, N.; Demon, S.Z.N.; Jasmani, L.; Yunus, W.M.Z.W.; Yaacob, W.M.H.W.; Salleh, E.M.; Norizan, M.N.; et al. Comparative Study on of CellulExtraction ose Fiber from Rice Straw Waste from Chemo-Mechanical and Pulping Method. Polymers 2022, 14, 387. https://doi.org/10.3390/polym14030387

20. Satyanarayana KG, Flores-Sahagun THS, Santos LPD, et al., 2013. Characterization of blue agave bagasse fibers of Mexico. Compos Part A: Appl Sci Manuf, 45:153–161. https://doi.org/10.1016/j.compositesa.2012.09.001

21. Simberloff, D. (2000). Global climate change and introduced species in United States forests. Science of the total environment, 262(3), 253–261.

22. Sindhu, R., Binod, P. & Pandey, A. (2015). Microbial Poly-3-Hydroxybutyrate and Related Copolymers. Industrial Biorefineries & white Biotechnology (pp-575–605).

23. Song, K., Zhu, X., Zhu, W. et al. Preparation and characterization of cellulose nanocrystal extracted from *Calotropis procera* biomass. Bioresour. Bioprocess. 6, 45 (2019). https://doi.org/10.1186/s40643-019-0279-z

24. Subedi, R.C., Bhuju, R.D., Bhatta, D.G., Pant, R.R., (2018), Climatic Variability and livelihood of rural farmers in Chisapani, Ramechhap, Nepal Journal of Environmental Science, 6, 47–59(2018).

25. Sun DY, Onyianta AJ, O’Rourke D, et al., 2020. A process for 460 J Zhejiang Univ-Sci B (Biomed & Biotechnol) 2021 22(6):450-461 | deriving high quality cellulose nanofibrils from water hyacinth invasive species. Cellulose, 27(7):3727–3740. https://doi.org/10.1007/s10570-020-03038-4

26. Vatseldutt, A. S. (2014).Isolation and Characterization of Polythene Degrading Bacteria from Polythene Dumped Garbage. International Journal of Pharmacy, 25(2), 205–206.

27. Vitousek, P. M., D’Antonio, C. M., Loope, L. L., Westbroo Mack, R., Simberloff, D., Lonsdale, M., Evans, H., Clout, M., & Bazzaz, F. (2000). Biotic invasions: cause, epidemiology, global consequences, and control. Ecological Applications, 10:689–710.

28. Yaowen, L., Ahmed, S., Sameen, D.E., Wang, Y., Rui, L., Dai, J., Suqing, L., Wen, Q. (2021). A review of cellulose and its derivatives in biopolymer-based for food packaging application. Trends in Food Science & Technology, 112, 532–546 (2021).

29. Wang, F., Zhang, Q., Huang, K., Li, J., Wang, K., Zhang, K., et al. (2020). Preparation and characterization of carboxymethyl cellulose containing quaternized chitosan for potential drug scarrier. International Journal of Biological Macromolecules, 154, 1392–1399 (2020).

